# Ligify: Automated genome mining for ligand-inducible transcription factors

**DOI:** 10.1101/2024.02.20.581298

**Authors:** Simon d’Oelsnitz, Andrew D. Ellington, David J. Ross

## Abstract

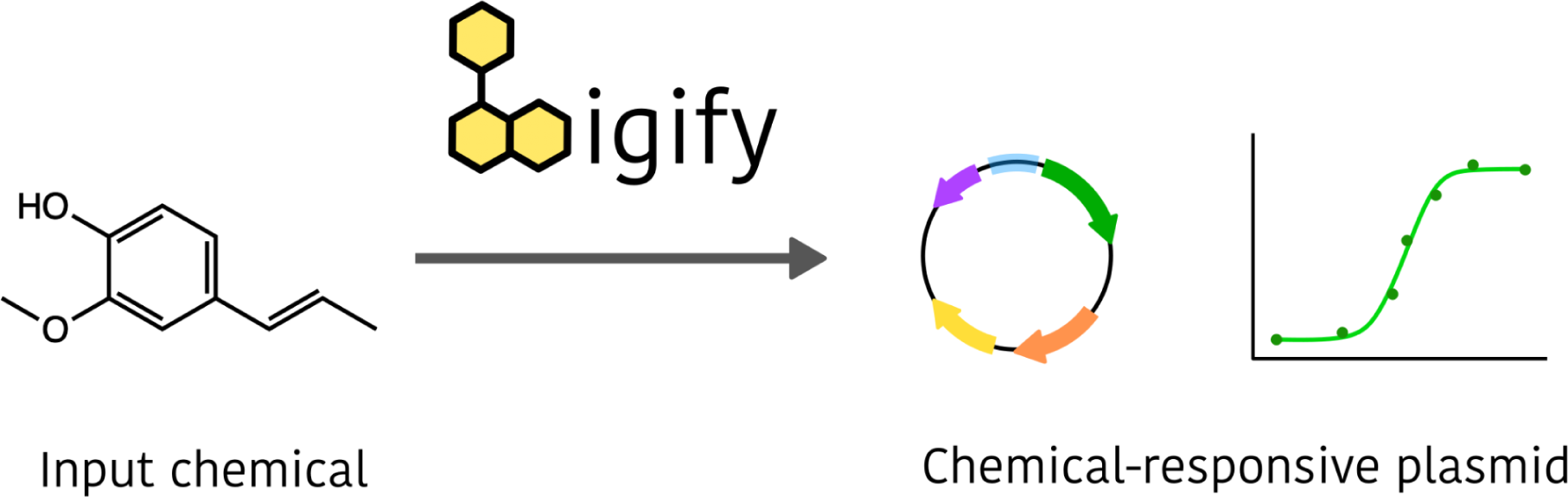

Prokaryotic transcription factors can be repurposed into biosensors for the ligand-inducible control of gene expression, but the landscape of chemical ligands for which biosensors exist is extremely limited. To expand this landscape, we developed Ligify, a web application that leverages information in enzyme reaction databases to predict transcription factors that may be responsive to user-defined chemicals. Candidate transcription factors are then incorporated into automatically generated plasmid sequences that are designed to express GFP in response to the target chemical. Our benchmarking analyses demonstrated that Ligify correctly predicted 31/100 previously validated biosensors, and highlighted strategies for further improvement. We then used Ligify to build a panel of genetic circuits that could induce a 47-fold, 5-fold, 9-fold, and 27-fold change in fluorescence in response to D-ribose, L-sorbose, isoeugenol, and 4-vinylphenol, respectively. Ligify should enhance the ability of researchers to quickly develop biosensors for an expanded range of chemicals, and is publicly available at https://ligify.streamlit.app.

## Introduction

Over the past couple decades, ligand-inducible prokaryotic transcription factors, herein called “regulators”, have become indispensable tools for chemical control of gene expression. Achieving higher dynamic ranges than riboswitches and being easier to manipulate than multi-component signaling cascades, regulators have seen increased use as biosensor components for biotechnology applications in various organisms, ranging from non-model bacterium^1^ to monkeys^2^. While early applications focused on simply inducing gene expression^3,4^, more emphasis has recently been given to the nature of the chemical inducer itself, which has enabled applications in high-throughput screening of enzymes^5^ and strains^6^, metabolic regulation^7^, production phenotype stabilization^8^, diagnostics^9^, and real-time measurement of biomarkers^10^.

To expand the bioengineer’s palette of available biosensors, regulators are often mined from bacterial genomes and “domesticated” into biosensors, which requires knowledge of (1) the regulator protein sequence, (2) the regulator’s cognate DNA binding sequence, or operator, and (3) the regulator’s cognate ligand. Domestication involves expressing the regulator on a plasmid alongside a reporter gene, often GFP, that is separately expressed using a promoter containing the regulator’s operator. The resulting plasmid is then tested for ligand-induced expression of the reporter gene. This general workflow has been used to create biosensors for terpenes^11^, cannabinoids^12^, sugars^13^, and flavonoids^14^, among others. Furthermore, advances in DNA synthesis and automated workflows have enabled the high-throughput mining and domestication of large panels of regulators into biosensors for various metabolites^15^.

While effective, genome mining is a limited and oftentimes laborious process. Identifying a candidate regulator for a chemical of interest typically requires manually searching for literature reports that characterize a microbial enzyme acting on the target chemical, information which is typically restricted to well studied organisms. Alternatively, growing genome databases represent a treasure trove of largely unexplored regulators. Leveraging these rich datasets, Hanko et al created a bioinformatic tool, TFBMiner, that partially automates the genome mining workflow^16^. The program takes a chemical input, which it uses to search for associated enzymes in the KEGG database^17^, and subsequently returns neighboring regulators as biosensor candidates. However, TFBMiner suffers from several notable drawbacks, including the requirement for the regulator to be divergently expressed from the chemical-associated enzyme, the redundancy of highly similar candidate regulators, and poor accessibility of the command line interface to less computationally proficient biologists.

To facilitate the mining and domestication of regulators for bioengineering applications, we developed Ligify, a fully automated chemical-to-plasmid program accessible via a user-friendly web application. We benchmarked Ligify against a curated dataset of 100 previously validated regulator:ligand pairs, revealing that Ligify correctly predicted a greater number of regulators than TFBMiner (31 vs 26, respectively), while also highlighting strategies for further improvement. We then experimentally validated Ligify-generated plasmids in *E. coli*, resulting in four novel biosensors for D-Ribose, L-Sorbose, Isoeugenol, and 4-Vinylphenol with induction ratios up to 47-fold. We expect Ligify to make regulator biosensor development simpler and more accessible to the broader bioengineering community.

## Results

### Workflow for biosensor prediction

Two key assumptions were made in designing a program that would accept a small molecule input and return a candidate biosensor: first, that regulators responsive to a small molecule are co-expressed with neighboring enzymes that act on that small molecule; and second, that a regulator should control its own expression. While these two assumptions are not always true in nature^18^, they enable a systematic method by which novel transcription factors could be computationally identified.

The Ligify workflow starts by using the small molecule input (in SMILES, text, or molecular drawing format), to fetch enzymatic reaction IDs associated with that small molecule from the Rhea database^19^ (**Figure 1**). Reaction IDs are then used to fetch all bacterial enzymes in Uniprot associated with each reaction, and highly homologous enzymes are filtered to limit redundancy. For each unique enzyme associated with the small molecule input, the enzyme’s respective operon is fetched from NCBI, which is scanned for genes annotated as “regulator”, “repressor”, or “activator”. Operons typically contain two divergently expressed sets of genes. If a regulator is found, the region of DNA between the two divergently expressed sets of genes is extracted; this region is hypothesized to contain a promoter responsive to the input small molecule. Finally, a rank for each regulator is calculated based on a set of heuristics, including the regulator-enzyme distance, the operon size, and the number of regulators in the operon. Specifically, a high rank is produced for regulators that are located nearby the small molecule-associated enzyme and that are within an operon that had few genes and only one regulator. A series of advanced options may be adjusted to speed up processing time by limiting the number of reactions, enzymes, and regulators that are evaluated. If no regulators are returned, Ligify will suggest alternative molecules similar to the input chemical that are also found in the Rhea database.

**Figure 1.**
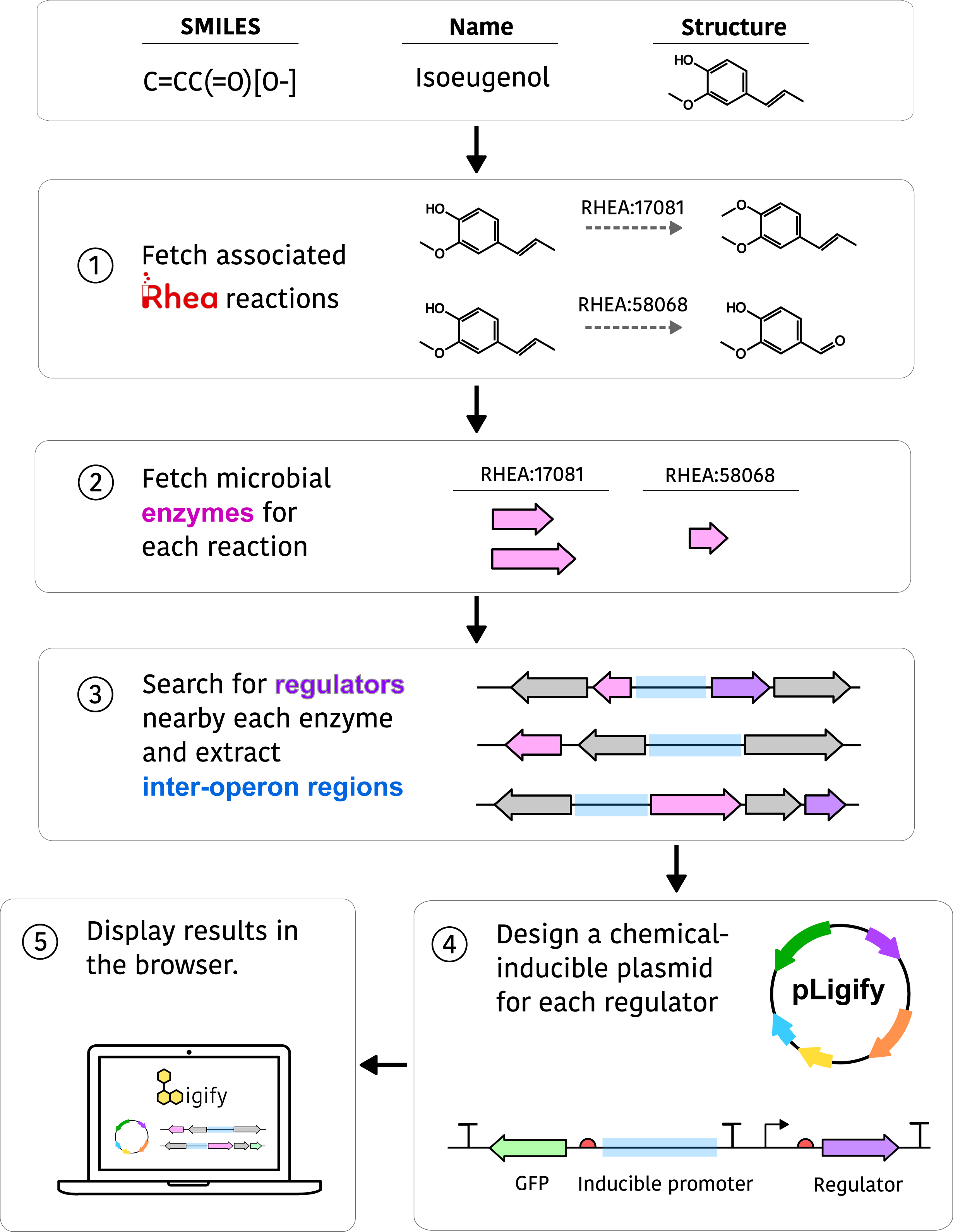
The Ligify workflow. An input chemical name, SMILES code, or structure is used to fetch reactions associated with that chemical in the Rhea database (1). Reactions associated with the input chemical are fetched from the Rhea database. (2) Phylogenetically divergent microbial enzymes associated with each reaction are returned. (3) The local genetic context of each enzyme is screened for regulators and if a regulator is identified, (4) a plasmid is designed wherein the regulator is expressed from a constitutive promoter and the GFP gene is expressed from the inter-operon region assumed to contain a regulated promoter. (5) Predicted regulators and associated metrics are displayed in a web browser.

To provide an actionable output for the user, a plasmid is generated for each predicted regulator that is designed to express GFP in response to the small molecule input. The plasmid contains a low copy p15A origin and expresses the regulator and GFP from two separate promoters. The regulator is expressed using a medium strength sigma-70 promoter^11^, the RiboJ ribozyme insulator^20^, and a bicistronic translational element^21^, to enable context-independent tuning of transcriptional and translational strengths (**Supplementary Figure 1**). GFP is expressed from a synthetic expression element consisting of the inter-operon region hypothesized to contain a promoter controlled by the predicted regulator, the ElvJ ribozyme, and a strong synthetic RBS.

Upon completion of the Ligify workflow, results are interactively displayed in a Web browser (**Supplementary Figure 2**). Search metrics describe how many reactions, enzymes, operons, and regulators were evaluated from the search criteria. For each predicted regulator, metadata is displayed on the regulator, its rank, the associated enzyme, the sequence of the operon, the sequence of the inter-operon region, and possible alternative inducer molecules based on reactions catalyzed by other enzymes in the operon. Additionally, a link is available to download the designed small molecule-responsive plasmid in GenBank format.

### Benchmarking Ligify

To benchmark the accuracy of Ligify, we curated a dataset of 100 previously validated biosensor:chemical interactions from the groov^DB^ database^22^, and other literature sources (**Supplementary Dataset 1**). Ligify was compared with the command-line tool TFBMiner, which to our knowledge is the only other biosensor search tool that uses comparable inputs and outputs^16^. Similar to Ligify, TFBMiner uses a guilt-by-association model to predict transcription factor specificity via proximal enzyme specificity. However, in contrast to Ligify, TFBMiner queries the Kyoto Encyclopedia of Genes and Genomes (KEGG) database for information on protein-ligand associations. Furthermore, TFBMiner uses a different search algorithm that only extracts regulators expressed divergently from chemical-associated enzymes, whereas Ligify extracts regulators expressed in both the same and opposite directions as enzymes. Each small molecule in our dataset was used as an input for both Ligify and TFBMiner, and the predicted regulators returned by each model were compared to the corresponding experimentally verified regulators in the dataset (**Figure 2a)**. If the verified regulator was found in the list of candidate regulators returned by the tool, the prediction was counted as a success. Overall, Ligify correctly predicted a greater number of regulators than TFBMiner; Ligify predicted 31/100 regulators and TFBMiner predicted 26/100 regulators (**Figure 2b**).

**Figure 2.**
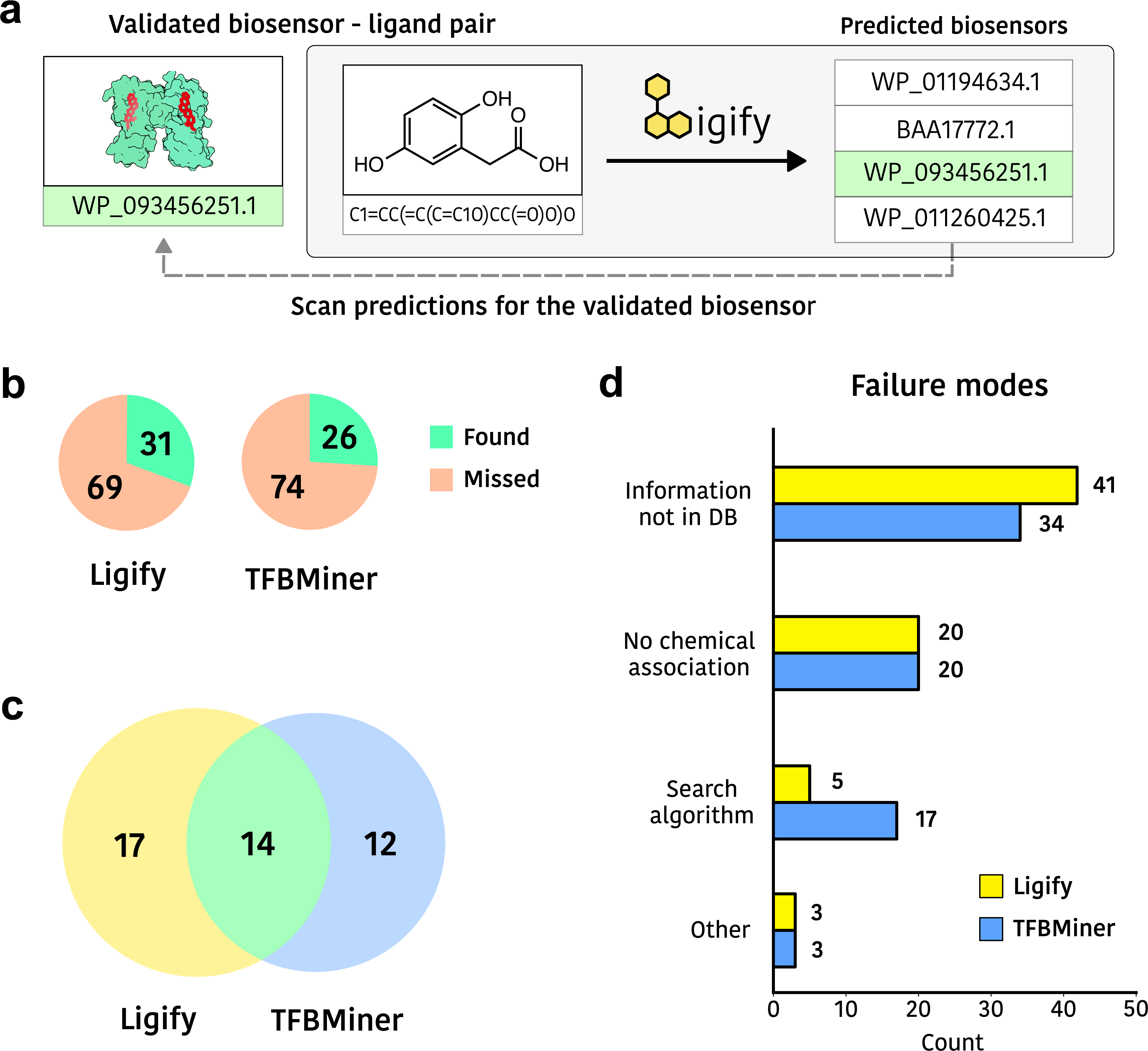
Benchmarking Ligify. (a) Benchmarking workflow. Ligands from a dataset of experimentally validated ligand-regulator pairs are passed into Ligify, and the predicted regulators are then compared to the sequence of the validated regulator. (b) The accuracy of Ligify and TFBMiner. (c) Comparison of the regulators correctly predicted by Ligify and TFBMiner. (d) Analysis of the failure modes encountered during benchmarking of Ligify and TFBMiner. An in depth analysis of failure modes, as well as all associated benchmarking metrics, can be found in **Supplementary Data 1**.

Interestingly, only 14 regulators were predicted by both models, while 17 or 12 regulators were uniquely predicted by Ligify or TFBMiner, respectively, suggesting that each model has different informatic strengths (**Figure 2c**). To further understand the differences in predictions made by Ligify and TFBMiner, we grouped failure modes into four separate categories: (1) a reaction is associated with the small molecule, but this association is not documented in the queried reaction database (2), the search algorithm could not identify the regulator (3), no reactions are associated with the input small molecule and (4) all other failure modes.

A greater number of failed predictions were attributed to insufficient information in the reaction database for Ligify than for TFBMiner (**Figure 2d**). This can be explained by the fact that each tool queries different reaction databases; TFBMiner queries KEGG while Ligify queries Rhea. In contrast, TFBMiner failed to predict 17 regulators based on the regulator search algorithm, while Ligify missed five regulators on this basis. This difference can be explained by the fact that TFBMiner uses a restrictive search algorithm that can only identify regulators that are expressed in the opposite direction as the small molecule-associated enzyme, while Ligify can identify regulators expressed in both the same and opposite directions as the enzyme. Indeed, 13 regulators missed by TFBMiner were expressed convergently relative to the small molecule-associated enzyme (**Supplementary Table 1**).

Both models failed to predict the same 20 regulators for small molecules that were not associated with any enzymatic reactions. Some of these regulators are expressed divergently from transporters or uncharacterized proteins, and others are thought to regulate promoters outside their local genetic context (see Supplementary Dataset 1). In addition, both models failed to predict three regulators for reasons such as the validated inducer molecule being a chemical analog of a natural metabolite or the small molecule-associated enzyme promiscuously acting on many small molecules.

### Using Ligify to identify novel biosensors

To demonstrate the utility of Ligify for biosensor discovery, we predicted and then experimentally tested several transcriptional regulators. We targeted sugars, volatiles, and neurochemicals as inducer molecules, since they are generally difficult to measure via traditional analytical approaches and are valuable for applications in pharmaceutical screening, cell-cell communication, and diagnostic applications. The chemicals D-Ribose, D-Arabitol, L-Sorbose, 4-Vinylphenol, Guaethol, Isoeugenol, Dopamine, and L-Carnitine were passed into Ligify as input molecules in SMILES format. These queries returned several regulators from the LacI, DeoR, LysR, AraC, and LuxR structural families (**Supplementary Table 2**) that were used to auto-generate biosensor plasmids in GenBank format (see **Supplementary Data 2** and **Figure 1**). The regulators returned for Dopamine (dadR) and Isoeugenol (iemR) had been previously identified and partially characterized^23,24^, but the remaining regulators had not been studied. For all ligands except for D-Ribose, one biosensor plasmid was designed and synthesized that contained the predicted regulator with the highest rank, and its associated promoter. For D-Ribose, four biosensor plasmids expressing regulators from divergent bacterial hosts were cloned to test the ability of Ligify to identify highly dissimilar regulators that bind to the same ligand. Each regulator for D-Ribose had between 29 % and 52% sequence similarity to one another (**Supplementary Table 3**).

Upon induction of cells containing biosensor plasmids with their cognate ligand, 7 of the 11 strains produced a significant fluorescent response (**Figure 3a**). The regulators sorR, vprR, and iemR all responded to their cognate ligand, while the regulators dalR, gcoR, dadR, and cdhR showed no response. Encouragingly, all four predicted D-Ribose regulators were found to respond to D-Ribose, despite having divergent sequences, with induction ratios ranging between 2-fold to 20-fold. These results indicate that Ligify can facilitate the identification of functionally homologous regulatory operons in divergent bacterial species.

**Figure 3.**
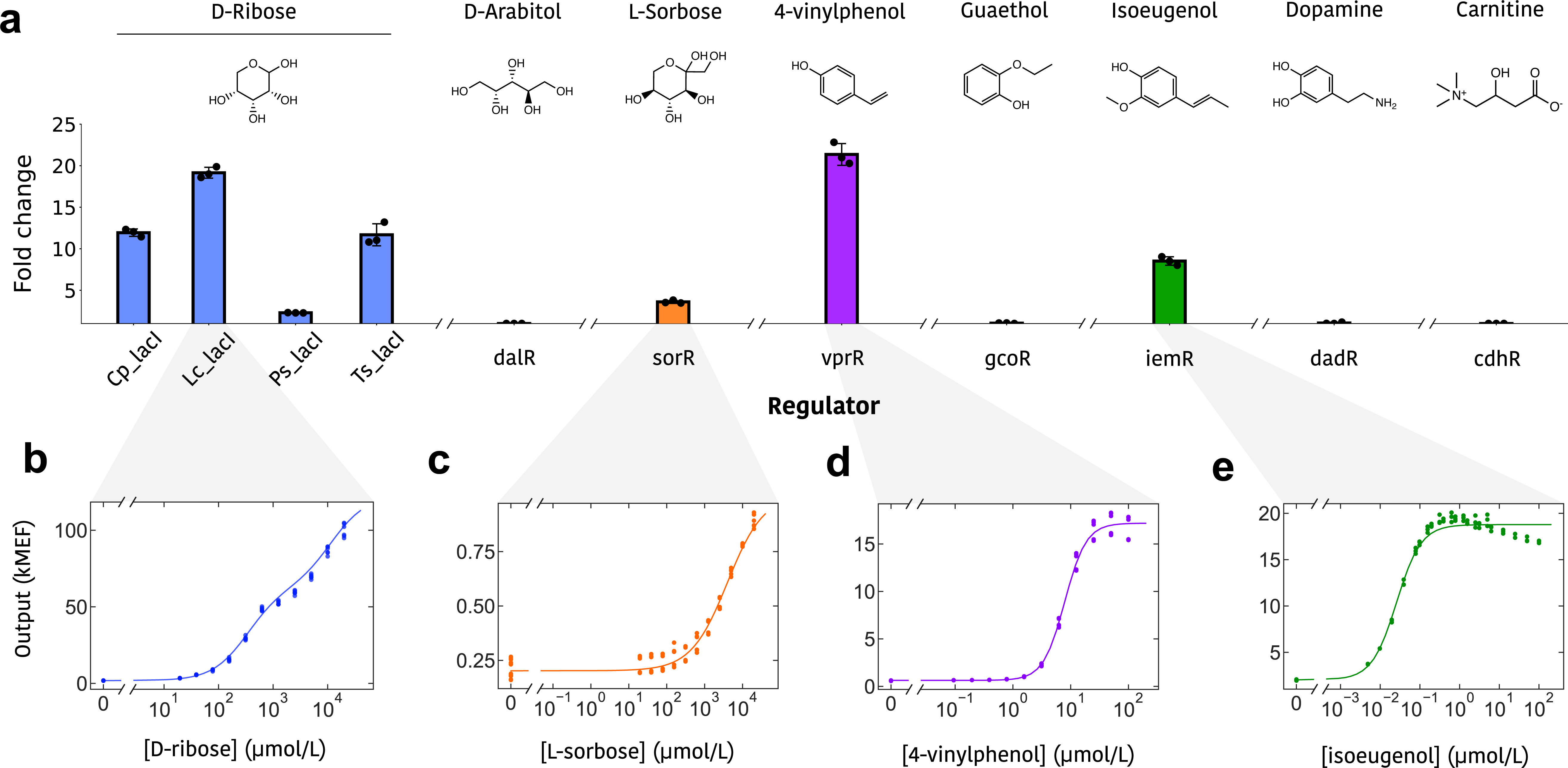
Validation of auto-generated reporter plasmids. (a) The fluorescent response of *E. coli* bearing Ligify-generated plasmid designs when induced with predicted effector molecules. The ligand concentration for induction was 5 mM for D-Ribose and L-Sorbose and 1 mM for all other ligands, which was chosen based on the compound’s solubility limit in 1% DMSO. Error bars represent the standard deviation from the mean. (b-e) Dose response measurements of the LcLacI, sorR, vprR, and iemR regulators to their cognate ligands, respectively. Assays were performed in biological triplicate and individual data points are shown.

To confirm the dose-dependent response of regulators to their cognate ligands, we measured the fluorescence of cells containing the LcLacI, SorR, VprR, and IemR regulators with varying concentrations of D-Ribose, L-Sorbose, 4-Vinylphenol, and Isoeugenol, respectively. An expected sigmoidal response was produced for the SorR, VprR, and IemR regulators (**Figure 3b-e**). Interestingly, the IemR regulator was found to be extremely sensitive to Isoeugenol, with an EC_50_ of nearly 25 nmol/L. The LcLacI-bearing strain produced a dose response that fit a double-sigmoid function, which may be due to the catabolism of D-Ribose by *E. coli*. These results demonstrate that Ligify is capable of creating functional biosensor plasmids for previously identified regulators (IemR), as well as for completely uncharacterized regulators, including SorR, VprR, and the four D-Ribose regulators.

### Characterization of biosensor selectivity

One key advantage of transcription factors is their capacity for exquisite chemical specificity. Some transcription factors have been reported to precisely distinguish between particular regioisomers or stereoisomers of a chemical scaffold, which would traditionally require lengthy chromatographic separation^25,26^. Conversely, non-specific transcription factors that promiscuously respond to several small molecules have been reported to be excellent starting points for evolution towards specific biosensors^27^. Thus, being able to utilize Ligify-generated circuits for the rapid characterization of specificity is a key feature of its incorporation into a variety of workflows.

To better understand the selectivity of the LcLacI, SorR, VprR, and IemR regulators, we characterized their responses to three analogs of their cognate inducer molecule. The LcLacI regulator displayed a highly selective response to D-Ribose, and responded poorly to the stereoisomer L-Ribose and the two other five carbon sugars D-Arabitol and L-Arabitol (**Figure 4a**). VprR also demonstrated extremely high selectivity, yielding a 27-fold response to 100 μmol/L of its cognate ligand, 4-vinylphenol, and a 2.6-fold response to 1000 μmol/L of the highly similar analog 4-ethylphenol (**Figure 4b**, **Table 1**). The SorR regulator displayed a higher selectivity for L-Sorbose (5.1-fold response) relative to D-Sorbose (1.5-fold response), but also appeared to respond to the six carbon sugar analogs D-Glucose and D-Galactose (**Figure 4c**, **Table 1**). IemR produced a response to all analogs tested, with sensitivities ranked highest to lowest as follows: isoeugenol (25 nmol/L), 4-vinylphenol (610 nmol/L), eugenol (65 μmol/L), 4-ethylguaiacol (207 μmol/L) (**Figure 4d**, **Table 1**). These data agree with a previous study on IemR selectivity that suggested that the propenyl group of isoeugenol was more critical for binding and activating IemR than the methoxy group^24^. Our flow cytometry analysis indicated that cell fluorescence distributions were unimodal (as expected) in all regulator-chemical induction conditions, suggesting that induction was not significantly limited by chemical transport, as reported for other biosensors^28^ (**Supplementary Figure 3**). Thus, the Ligify-designed circuits were able to quickly characterize the identified sensors as either being highly specific (LcLacI and VprR), and thus potentially useful for integration into metabolic control circuits, or semi-specific (SorR and IemR), and thus of potential utility for further engineering towards a number of more unique specificities.

**Figure 4.**
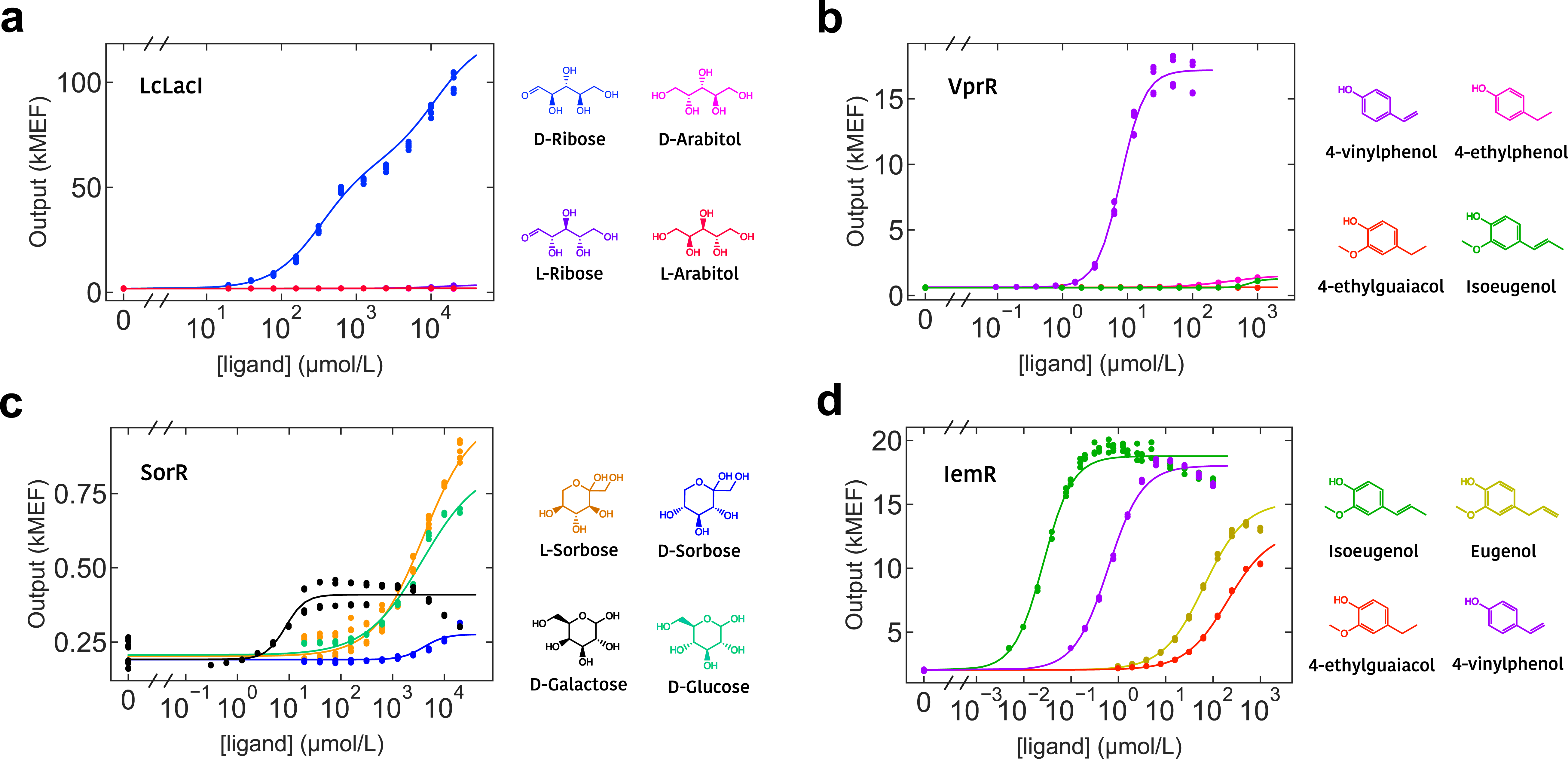
Characterization of biosensor selectivity. (a-d) Dose response of Ligify-generated plasmids to the cognate ligand and ligand analogs for the LcLacI, vprR, sorR, and iemR regulators, respectively. Assays were performed in at least biological triplicate and individual data points are shown. The maximum ligand concentration was chosen based on the compound’s solubility limit in 1% DMSO. Associated performance metrics are summarized in **Table 1**.

**Table 1.**
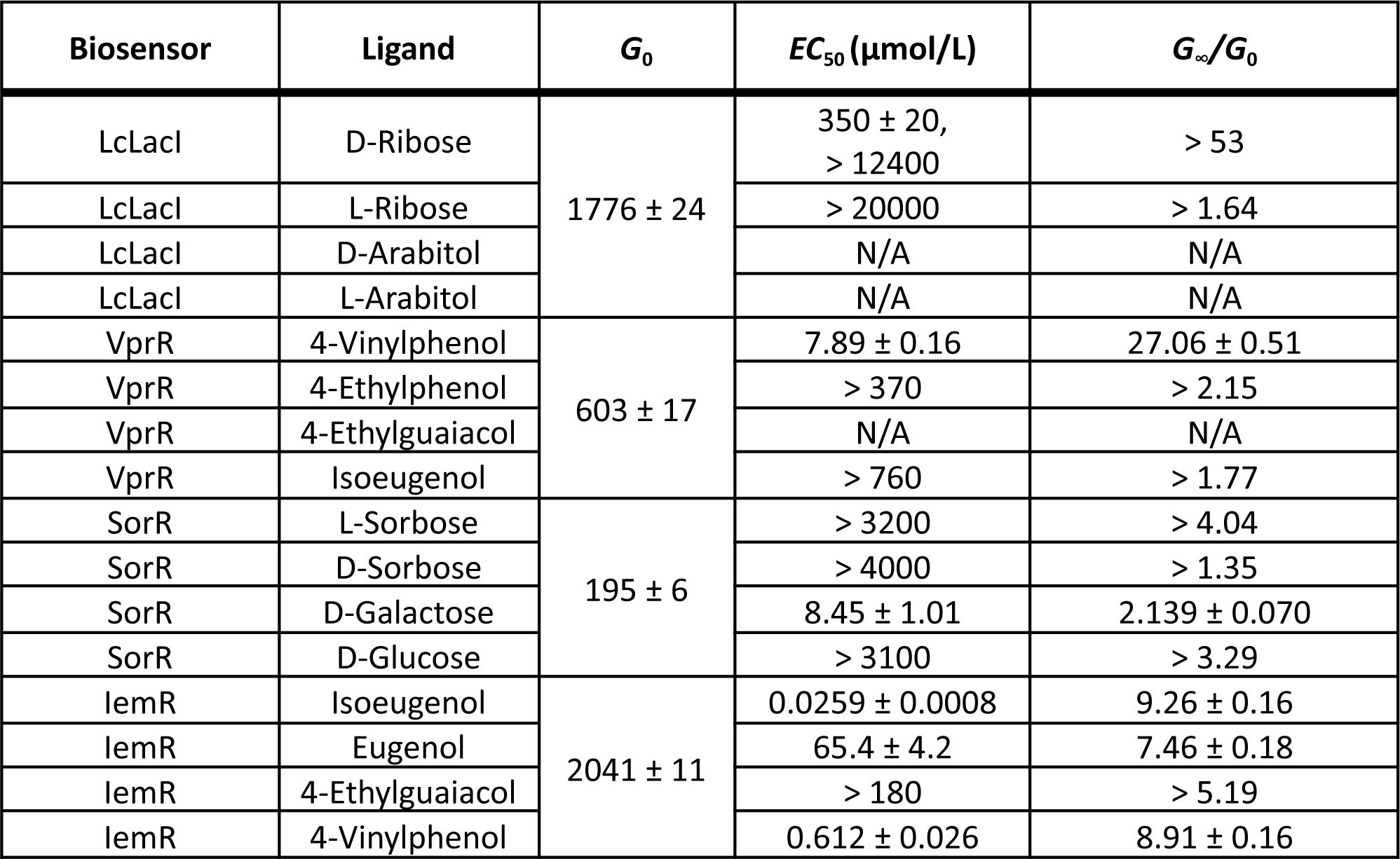
Performance metrics of characterized genetic biosensors. *G*_0_ values are reported as molecules of equivalent fluorescein. *G*_0_ represents the response without any ligand supplemented, and *G*_∞_/*G*_0_ represents the fold change in fluorescence with versus without the ligand added. For responses where the *EC*_50_ is close to or greater than the maximum concentration tested, the 95% confidence lower bound for the *EC*_50_ and the *G,*/*G,* ratio is reported. For regulator-ligand combinations that did not show any significant response, “N/A’’ is reported. Two apparent *EC*_50_s are reported for the response of LcLacI to D-Ribose.

| Biosensor | Ligand | $G_0$ | $EC_{50}$ ($\mu\text{mol/L}$) | $G_{\infty}/G_0$ |
| --- | --- | --- | --- | --- |
| LcLacI | D-Ribose | $1776 \pm 24$ | $350 \pm 20$,<br>> 12400 | > 53 |
| LcLacI | L-Ribose |  | > 20000 | > 1.64 |
| LcLacI | D-Arabitol |  | N/A | N/A |
| LcLacI | L-Arabitol |  | N/A | N/A |
| VprR | 4-Vinylphenol | $603 \pm 17$ | $7.89 \pm 0.16$ | $27.06 \pm 0.51$ |
| VprR | 4-Ethylphenol |  | > 370 | > 2.15 |
| VprR | 4-Ethylguaiaicol |  | N/A | N/A |
| VprR | Isoeugenol |  | > 760 | > 1.77 |
| SorR | L-Sorbose | $195 \pm 6$ | > 3200 | > 4.04 |
| SorR | D-Sorbose |  | > 4000 | > 1.35 |
| SorR | D-Galactose | | $8.45 \pm 1.01$ | $2.139 \pm 0.070$ |
| SorR | D-Glucose |  | > 3100 | > 3.29 |
| IemR | Isoeugenol | $2041 \pm 11$ | $0.0259 \pm 0.0008$ | $9.26 \pm 0.16$ |
| IemR | Eugenol | | $65.4 \pm 4.2$ | $7.46 \pm 0.18$ |
| IemR | 4-Ethylguaiaicol |  | > 180 | > 5.19 |
| IemR | 4-Vinylphenol | | $0.612 \pm 0.026$ | $8.91 \pm 0.16$ |

## Discussion

Ligify is a web-accessible bioinformatic tool that fully automates the process of discovering and designing biosensor circuits for user-defined small molecules by extracting functional elements from bacterial genomes. Ligify was able to correctly predict transcription factors for 31 of 100 previously validated transcription factor:ligand pairs across diverse protein structural families, which now represents state-of-the-art performance for this prediction task. In consequence, Ligify is poised to circumvent the slow manual process of mining transcription factors from bacterial genomes, and accelerate the discovery of genetic sensors for synthetic biology and biotechnology applications.

Our benchmarking results revealed strategies to further improve the model’s accuracy. One of the largest failure modes was caused by the lack of a small molecule-enzyme association in the queried reaction database. Interestingly, TFBMiner was able to correctly predict regulators missed by Ligify for this failure mode, and vice versa, largely because TFBMiner queries KEGG while Ligify queries Rhea. This suggests that querying both KEGG and Rhea, along with other reaction databases such as BRENDA^29^ or Expasy^30^, may provide additional access to enzyme-ligand pairs, further improving accuracy. A second addressable failure mode is caused by the algorithm used to search for regulators within an operon. Five regulators were missed by Ligify for this reason, largely due to issues with fetching the genetic context or the exclusion of the regulator gene from the calculated operon (**Supplementary Table 4**). Updating the regulator search algorithm to capture these missed regulators would provide another mechanism to increase the accuracy of Ligify.

Functionally, seven of eleven plasmid designs generated *de novo* by Ligify were shown to produce a fluorescent response to the intended small molecule in *E. coli*. This result is surprising, given that the associated regulators were sourced from diverse organisms, belong to distinct structural families, and that a generic plasmid template was used for all regulators without promoter or RBS tuning, which is sometimes required to produce a functional design^13^. On average, biosensors that produced a functional response generally received a higher score via our ranking system than their unresponsive counterparts, indicating that the rank returned by Ligify may serve as a useful guide for decision making (**Supplementary Table 2**). Further analysis of the regulators that did not produce a response highlight possible failure mechanisms (**Figure 3a**). In previous work, the enzyme associated with gcoR was characterized to promiscuously react with various ligands, suggesting that an analog of guaethol may be the true inducer molecule^31^. The dadR regulator was previously shown to respond to dopamine in its native organism, although it is a transmembrane protein that may require a partner protein to mediate a transcriptional response^23^. Finally, the hypothesized ligand of dalR, D-Arabitol, may require the expression of a transport pump for import, as is typical for polar sugars^32^. These observations may help develop a more informative ranking system. For example, regulators that contain a transmembrane domain or that are associated with a promiscuous enzyme may receive a lower rank, while regulators that are homologous to experimentally validated biosensors may receive a higher rank.

Ultimately, without any manual intervention, Ligify was able to design plasmids that were responsive to D-Ribose, L-Sorbose, Isoeugenol, and 4-Vinylphenol with induction ratios of 47-fold, 5-fold, 9-fold, and 27-fold, respectively. Given that each plasmid incorporated a transcription factor belonging to a distinct family (LaLacI: LacI, SorR: DeoR, IemR: AraC, VprR: LysR), Ligify-generated designs appear to be able to generalize to almost any type of transcription factor. Furthermore, using Ligify, we identified and validated four D-Ribose-responsive regulators from divergent species, suggesting that Ligify might prove capable of mapping regulator orthologs across diverse prokaryotic phylogenies.

In the future, Ligify and its derivatives may play a key role in the overall goal of mapping the protein-ligand-DNA atlas of all prokaryotic ligand-inducible transcription factors. Predictions from Ligify can be used alongside operator prediction tools, such as Snowprint^33^, to narrow in on probable regions of DNA bound by the regulator. For example, we used Snowprint to isolate conserved inverted repeat sequences within promoters controlled by D-Ribose responsive transcription factors identified by Ligify, which now represent candidate operator sequences (**Supplementary Figure 4**). Ligify may also potentially be run “in reverse” to suggest possible effector ligands for an input regulator, using the same underlying algorithms. Data collected using Ligify and its derivatives may further be used to train machine learning models capable of predicting regulator:ligand pairings for regions of protein sequence space otherwise inaccessible to heuristic models. Existing protein-chemical machine learning models created by the drug discovery community, such as DeepDTA^34^, may already be suitable for this task.

In summary, Ligify is a generalizable ligand-to-regulator tool for biosensor discovery and design. Requiring just one input, the tool is easy to use and delivers an actionable output: a complete plasmid design that can be directly tested for response to the input chemical. Paired with the regulator-to-operator tool Snowprint^33^, and the groov^DB^ biosensor database^22^, Ligify fits within an ecosystem of web-based resources for the characterization and organization of genetic biosensors.

## Methods

### Benchmarking

Validated ligand-regulator pairs were collected from literature sources (**Supplementary Data 1**) and groov^DB^, a public database of ligand-inducible transcription factors^22^. Using this dataset, the SMILES code of each ligand was used as the input to a Ligify query to produce a set of predicted regulators and prediction metrics for each query (**Supplementary Data 1**). Notably, SMILES codes of biologically-relevant isomers were used for chemicals that have multiple isomers; For example, the SMILES code corresponding to the chemical “D-ribofuranose” was used instead of that for the chemical “D-ribose”. The protein sequences for the validated regulators and the predicted regulators (i.e., returned by Ligify) were retrieved using NCBI’s eFetch tool, and BLAST was used to compare the predicted regulator sequences to the validated regulator sequences. A prediction was considered correct if one of the predicted regulator sequences from a Ligify query exactly matched the corresponding validated regulator sequence.

### Strains, plasmids and media

*E. coli* DH10B (New England Biolabs) was used for all routine cloning and biosensor characterization. LB Miller (LB) medium (BD) was used for routine cloning, and M9 media was used for fluorescence assays, unless specifically noted. LB with 1.5% agar (BD) plates were used for routine cloning. M9 media was composed of the following filter-sterilized components: 33.9 g/L Disodium Phosphate, 15 g/L Monopotassium Phosphate, 2.5 g/L Sodium Chloride, 5 g/L Ammonium Chloride, 100 umol/L, CaCl_2_, 2 mmol/L MgSO_4_, 0.2% Casamino acids, 0.4% Glycerol, 50 ug/mL Kanamycin. The plasmids described in this work were constructed by Twist Biosciences. Schematic of the biosensor plasmid design used in this study is displayed in **Supplementary Figure 1**. Preparation of chemically competent cells was performed as follows. Briefly, 5 mL of an overnight culture of DH10B cells was mixed with 500 ml of LB and grown at 37 °C and 250 r.p.m. for three hours. Resulting cultures were centrifuged (3,500g, 4 °C, 10 min), and pellets were washed with ice cold 70 mL of chemical competence buffer (10% glycerol, 100 mM CaCl_2_) and centrifuged again (3,500g, 4 °C, 10 min). Pellets were then resuspended in 20 mL of ice cold chemical competence buffer. After 30 min on ice, cells were divided into 250 μL aliquots, flash frozen in liquid nitrogen, and stored at −80 °C until use. This method has been described previously^1^. For transformation, chemically competent cells were thawed on ice, then 20 μL of cells were mixed and incubated on ice with 1 μL of plasmid for 5 minutes. Then, cells were heat shocked at 42°C for 45 seconds, put back on ice for 2 minutes, and then 120 μL of SOC media was added. The resulting culture was shaken at 37 °C and 250 r.p.m. for 1 hour and 20 μL of this culture was then plated on an agar plate with appropriate antibiotics.

### Curation of biosensors for experimental validation

Target ligands were passed to Ligify in SMILES format. In cases where multiple regulator candidates were returned, two filters were applied to choose candidate regulators for testing. First, any literature on the associated reaction, returned by Ligify, was examined for cases whereby the ligand is shown to influence the expression of the associated operon. Second, the rank returned by Ligify was used to guide the decision. In the case of D-Ribose, four candidate regulators were chosen for validation.

### Chemicals

Sigma Aldrich was the vendor for the purchase of D-Ribose (R7500-5G), L-Arabitol (A3506-10G), L-Sorbose (85541-50G), D-Sorbose (S4887-100MG), 4-Ethylphenol (E44205-5G), 4-Ethylguaiacol (39774-1ML), Isoeugenol (I17206-5G), Eugenol (E51791-5G), Dopamine (H8502-5G), and Carnitine (8417740025). Fisher Scientific was the vendor for the purchase of L-Ribose (AC257910010), D-Arabitol (AC302870050), 4-Vinylphenol (AAL1090203), and Guaethol (AC370030050). D-Galactose was purchased from Fisher Scientific (part no. BP656). D-Glucose was purchased from Alfa Aesar (part no. A16828).

### Biosensor end-point assays

The following protocol was used to generate data shown in Figure 3a. Each biosensor plasmid was transformed into chemically competent E. coli cells. Three colonies were picked from each transformation and were grown overnight. The following day, 20 μl of each culture was then used to inoculate separate wells in a 2-ml 96-deep-well plate (Corning, P-DW-20-C-S) sealed with an AeraSeal film (Excel Scientific) containing 900 μl M9 media. After two hours of growth at 37 °C, cultures were induced with 100 µl M9 media containing either 10 μl solvent (either water or DMSO) or 100 μl M9 media containing the target compound dissolved in solvent (water or DMSO). Cultures were grown for an additional four hours at 37 °C and 1000 r.p.m. and subsequently centrifuged (3,500g, 4 °C, 10 min). Supernatant was removed, and cell pellets were resuspended in 1 ml PBS (137 mM NaCl, 2.7 mM KCl, 10 mM Na_2_HPO_4_, 1.8 mM KH_2_PO_4_, pH 7.4). One hundred microliters of the cell resuspension for each condition was transferred to a 96-well microtiter plate (Corning, 3904), from which the fluorescence (excitation, 485 nm; emission, 509 nm) and absorbance (600 nm) were measured using a plate reader (Biotek Neo2SM).

### Biosensor dose response assays

The following protocol was used to generate data shown in Figure 3b-e and Figure 4. Biosensor plasmids were transformed into *E. coli* DH10B cells, which were subsequently plated on LB agar plates containing appropriate antibiotics. Three individual colonies were picked for each transformation and were grown overnight in LB media. The following day, glycerol stocks of cultures were made by mixing 500 uL of culture with 500 uL of 40% glycerol, which were stored at −80°C. For measurement of the biosensor dose-response curves, cultures of *E. coli* containing each plasmid were started from glycerol stock scrapings in M9 media (5 mL culture in 15 mL snap-cap culture tubes). Cultures were grown overnight (16 h to 20 h) at 37°C with shaking at 300 rpm. Cultures were then loaded into a laboratory automation system for growth and measurement following a protocol similar to one previously described^2^. Briefly, that automated growth and measurement protocol was as follows:

1. Dilute cells 10-fold from snap-cap tubes into media in wells of a 96-well plate.
2. Plate type: Agilent part number 204799-100, square wells, 1 mL per well maximum volume
3. 0.5 mL per well culture volume
4. Apply a gas-permeable membrane to the 96-well plate.
5. Incubate for 12 h to 13.5 h in a plate reader (Biotek Neo2SM).
6. 37°C
7. Double-orbital shaking at 807 cycles per minute.
8. Measure optical density (OD600) and fluorescence (500 nm excitation and 530 nm emission) every 5 min during incubation, with continuous shaking between measurements.
9. Prepare a second 96-well plate with media and the addition of a dilution series of each ligand to be measured.
10. Ten minutes before the end of the incubation period, use a heated plate station to pre-warm the media in the second 96-well plate to 37°C.
11. At the end of the incubation period, remove the gas-permeable membrane from the first 96-well plate and dilute the cultures 50-fold from the first 96-well plate to the second 96-well plate.
12. Apply a gas-permeable membrane to the second 96-well plate.
13. Incubate for 3 h and 5 min in a plate reader (Biotek Neo2SM).
14. Same incubation and measurement settings as for the first incubation cycle.
15. Prepare a third 96-well plate with media and the addition of a dilution series of each ligand to be measured.
16. Ten minutes before the end of the second incubation period, use a heated plate station to pre-warm the media in the third 96-well plate to 37°C.
17. At the end of the incubation period, remove the gas-permeable membrane from the second

96-well plate and dilute the cultures 10-fold from the second 96-well plate to the third 96-well plate.

1. Apply a gas-permeable membrane to the third 96-well plate.
2. Incubate for 3 h and 5 min in a plate reader (Biotek Neo2SM).
3. Same incubation and measurement settings as for the first and second incubation cycles.
4. At the end of the incubation period, remove the gas-permeable membrane from the third 96-well plate and dilute the cultures 40-fold from the third 96-well plate into a flow cytometry sample plate.
5. Flow cytometry plate: round-bottom 96-well plate (Falcon part no. 351177).
6. μL of culture diluted into 195 μl of phosphate-buffered saline with 170 μg/mL chloramphenicol (Fisher BioReagents, cat. #BP904-100).

Samples were then measured using an Attune flow cytometer equipped with an autosampler using a 488 nm excitation laser and a 530 nm ± 15 nm bandpass emission filter.

An automated gating algorithm was used to discriminate cell events from non-cell events and singlet events from multi-plet events using blank samples that were measured with each batch of cell measurements^3^.

An additional automatic unsupervised gating procedure, similar to a previously described method^4^, was then used to select the region of the side-scatter vs. forward-scatter plots containing the highest density of singlet cell events. The additional automatic gating was set to retain 30% of the singlet cell events. Fluorescence calibration beads (Spherotech, part no. RCP-30-20A) were also measured with each flow cytometry plate to enable calibration of the flow cytometry data to molecules of equivalent fluorophore^5–7^.

### Statistics and Reproducibility

All data in the text are displayed as mean ± standard deviation unless specifically indicated. All experimental assays were performed in biological triplicate, which represent three individual bacterial colonies picked from an agar plate. Bar graphs were all plotted in Python 3.10.6 using Matplotlib 3.8. Dose-response curves were plotted in Python 3.11.4 using Matplotlib 3.7.2. Dose–response curves and EC_50_ values were estimated using cmdstanpy, version 1.1.0 (with Python 3.11.4).

### Building the Ligify web application

The Ligify web application was built in Python 3.10.8 using Streamlit 1.22.0 for both the frontend design and backend data architecture. During a complete prediction run, queries are made to the Rhea and NCBI databases. The Ligify web server is hosted on an AWS EC2 instance configured to use a custom domain name.

## Data availability

The relevant data are available from the corresponding authors upon request. NCBI RefSeq identifiers for all characterized regulators in this study can be found in **Supplementary Table 2.** NCBI RefSeq identifiers for all regulators used for Benchmarking can be found in **Supplementary Data 1**. Biosensor plasmids for the iemR, LcLacI, sorR, and vprR regulators were deposited in Addgene.

## Code availability

The source code and detailed instructions for use of Ligify are maintained in the GitHub repository located at https://github.com/simonsnitz/Ligify. Ligify is Open Access under an MIT License. Code used to generate bar plots presented in this manuscript is accessible in the GitHub repository located at https://github.com/simonsnitz/plotting. Code used for flow cytometry data analysis, to generate dose response functions and histograms is accessible in the GitHub repository here: https://github.com/djross22/ligafy_flow_analysis_git_repo.

## Acknowledgements

Funding from the National Institute of Standards and Technology (70NANB21H100) is acknowledged.

## Author Contributions

S.d’O. developed and benchmarked the Ligify software tool. S.d’O designed all experiments and S.d’O and D.J.R performed experiments and data analysis. The manuscript was written by S.d’O. with support from D. J. R.. A.D.E edited a final draft of the manuscript. S.d’O. supervised all aspects of the study.

