## Supplementary Information for "Ligify: Automated genome mining for ligand-inducible transcription factors"

Supplementary Figures 1 - 4

Supplementary Tables 1 - 4

#### Supplementary Figures

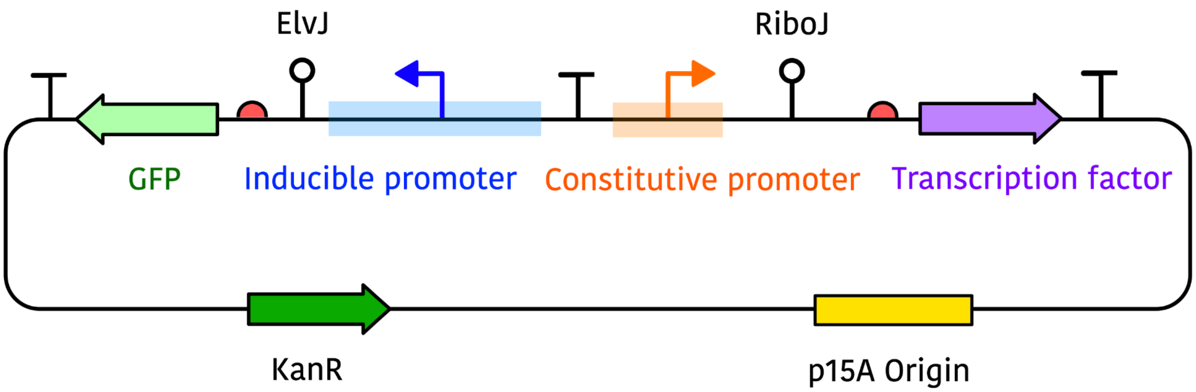


**Supplementary Figure 1:** Schematic of the auto-generated biosensor reporter design.
The transcription factor coding sequence (purple) and inducible promoter sequence (blue) are extracted from a microbial genome sequence. All other sequences are standardized in the pLigify design. Red half circles represent ribosome binding sequences and "T"s represent terminators. Please refer to pLigify plasmid GenBank files (**Supplementary Data 2**) for the complete sequence.

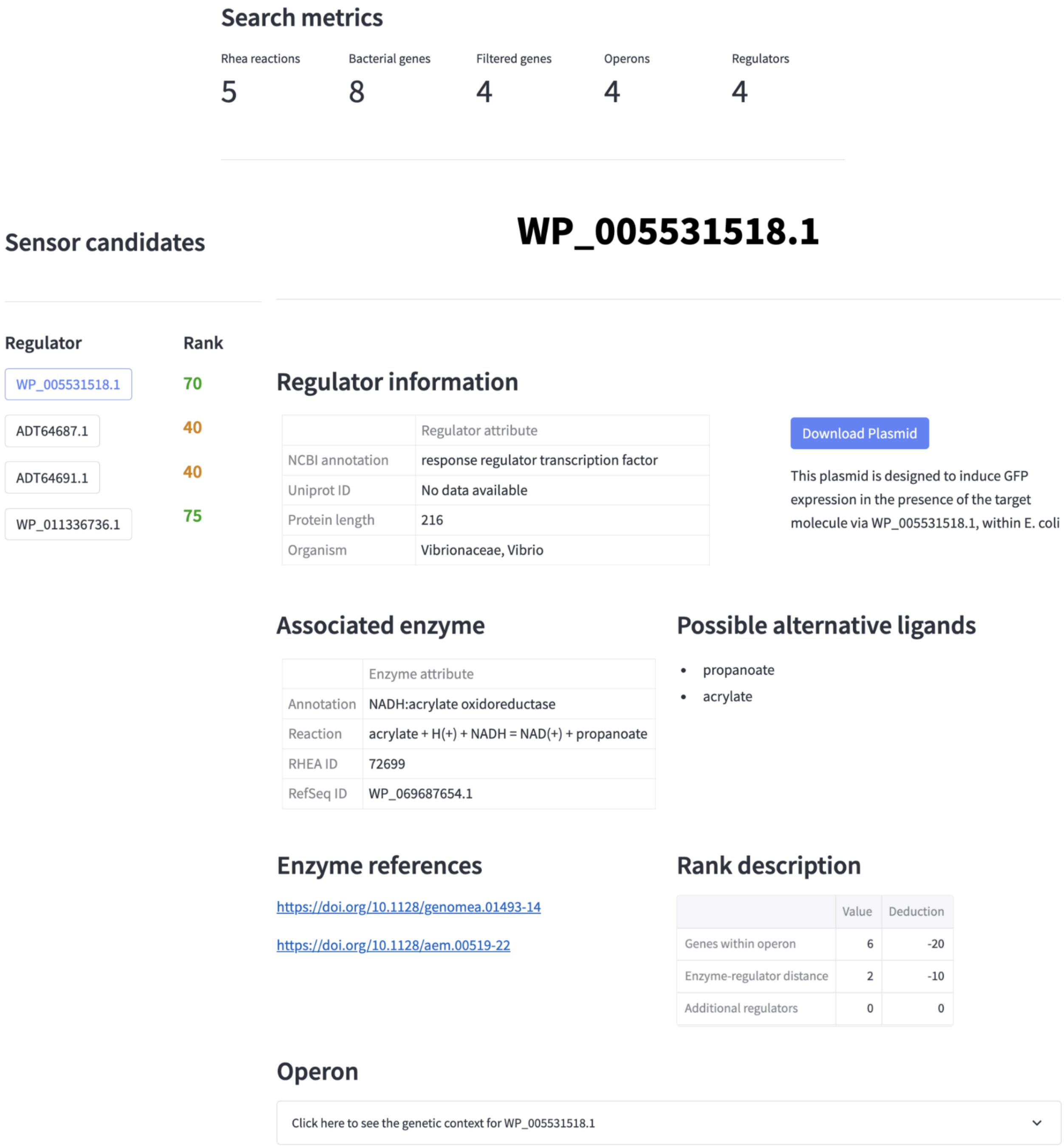


**Supplementary Figure 2:** Example of a biosensor report displayed in the Ligify web application

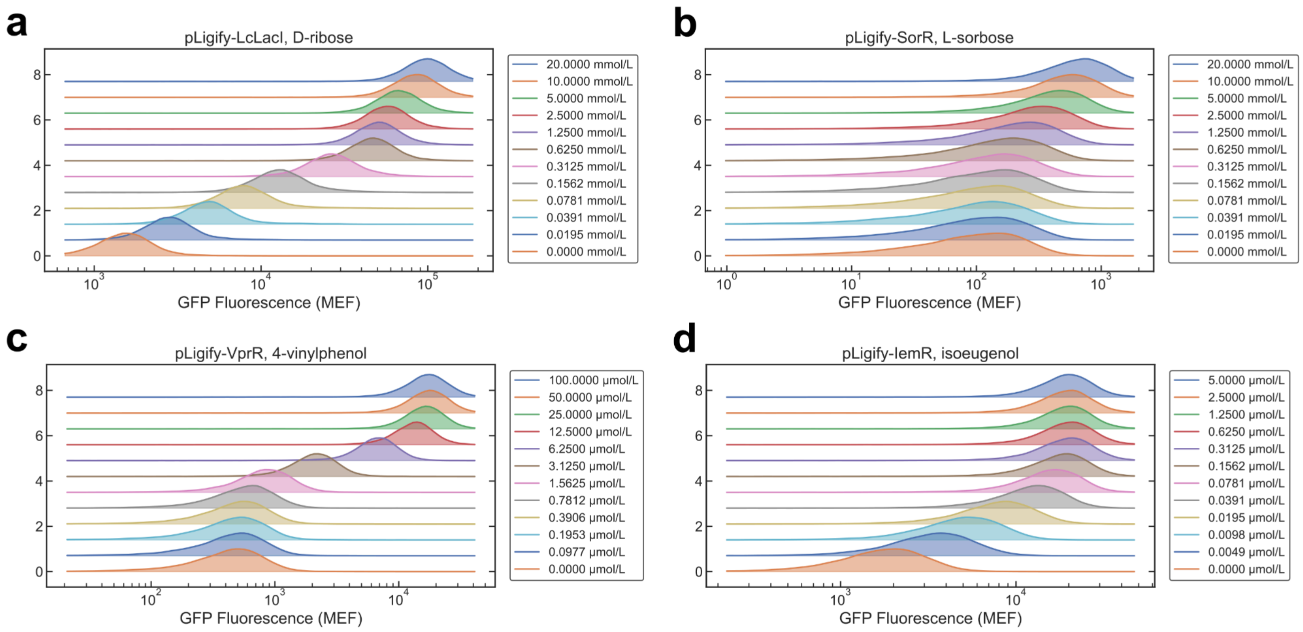


**Supplementary Figure 3:** Histograms of cell population distributions

(a-d) Distribution of cell fluorescence for *E. coli* cells bearing the LcLacI, sorR, vprR, or iemR biosensor plasmids upon induction with D-Ribose, L-Sorbose, 4-Vinylphenol, or Isoeugenol, respectively.

## WP_009051948.1

Conservation score

**63.945**

Sequences aligned

**28**

Predicted promoter region

AAATCAGGGCACAAAAAGCCCTGATTTTATAATAATTAAATCACGTGAAAAGTTATAAAAAATTTTTATTTTTATTATTGACAAAATGTCATT
TTTATTTTATTatataGTTAAGCGTTTTACTAAAACGTTAAACcaaatTTAATAGGTGATGAATTTA

Consensus sequence

**atttaGTTAAACGTTTTAGTAAAACGTTTAACtaaat**

Conservation motif logo

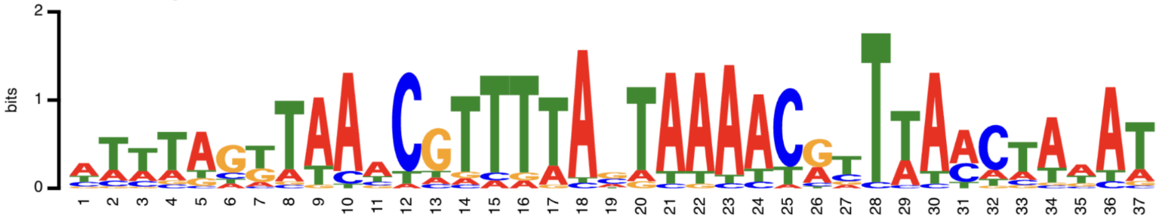

## WP_011834762.1

Conservation score

**95.407**

Sequences aligned

**27**

Predicted promoter region

ATTTAAAAGGCCTACTGAAAAGTAGGCTTTTTtgttaTTAAAACGAAATATAAAAAGGATTGACAAAATAAATGAAAACGGATACAATCATACA
TGTAAATGTGAGAACGTTCCCACAATTGAAAGGAGGAAGCCA

Consensus sequence

**AAAAGGCCTACTGAAAAGTAGGTTTTTTtgtta**

Conservation motif logo

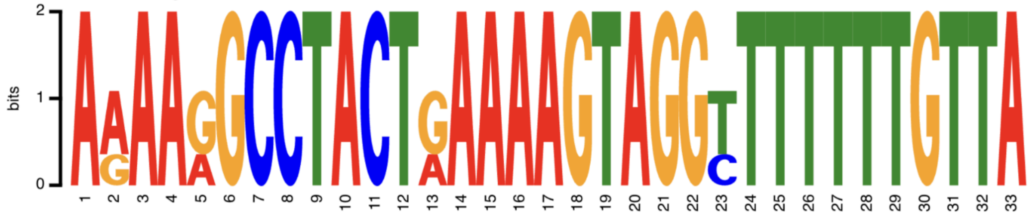

**Supplementary Figure 4:** Candidate operator sequences identified by Snowprint

### Supplementary Tables

| Regulator ID | Enzyme ID | Inducer | Operon |
|---|---|---|---|
| RSP_1435 | RSP_1433 | Acrylate | 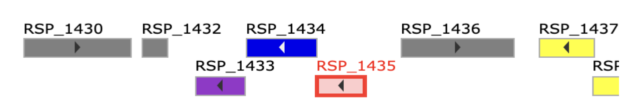 |
| JW0305 | JW0303 | Choline | 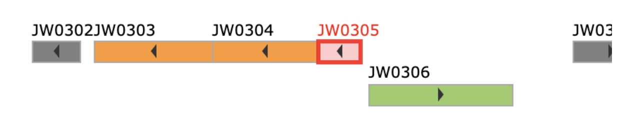 |
| JW3732 | JW3731 | D-Ribose | 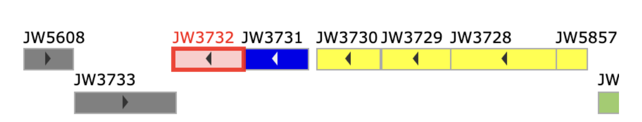 |
| JW0336 | JW0334 | Lactose | 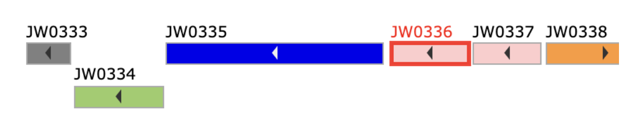 |
| JW5627 | JW5629 | D-Galactonate | 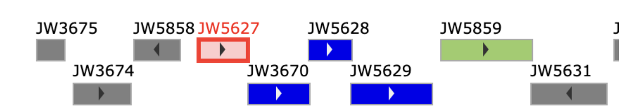 |
| BAB2_0369 | BAB2_0372 | Erythritol | 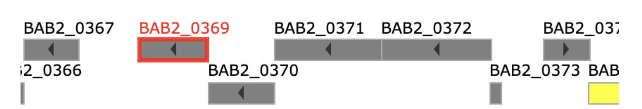 |
| BSU31210 | BSU31200 | L-Rhamnulose-1-phosphate | 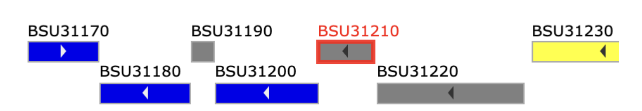 |
| JW0494 | JW0495 | Glyoxylate | 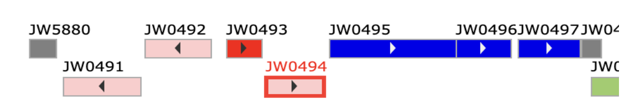 |
| JW0109 | JW0110 | Pyruvate | 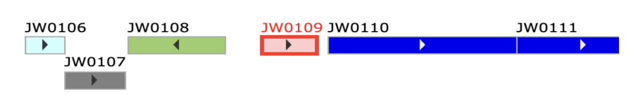 |
| H16_B0142 | H16_B0144 | (R)-acetoin | 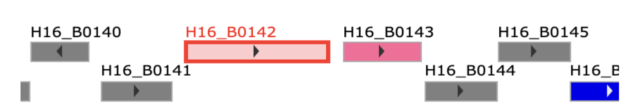 |
| AN232_30650 | AN232_30660 | Erythromycin A | 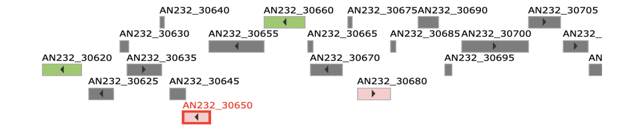 |
| BSU39990 | BSU39980 | Quercetin | 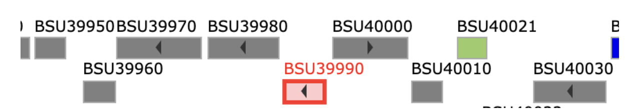 |
| PFLU_0359 | PFLU_0361 | Urocanate | 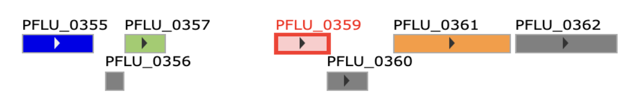 |

**Supplementary Table 1**: Convergently expressed regulators in the the benchmarking dataset

| Input SMILES | Ligand | Biosensor ID | Biosensor family | Associated enzyme | Rank |
|---|---|---|---|---|---|
| OC[C@H]1OC(O)[C@H](O)[C@@H]1O | D-Ribose | WP_012544132.1 | LacI | WP_012544127.1 | 70 |
| OC[C@H]1OC(O)[C@H](O)[C@@H]1O | D-Ribose | WP_010953404.1 | LacI | WP_010953405.1 | 70 |
| OC[C@H]1OC(O)[C@H](O)[C@@H]1O | D-Ribose | WP_009051948.1 | LacI | WP_009051947.1 | 55 |
| OC[C@H]1OC(O)[C@H](O)[C@@H]1O | D-Ribose | WP_011834762.1 | LacI | WP_011834763.1 | 15 |
| OC[C@@H](O)C(O)[C@H](O)CO | D-Arabitol | WP_011002058.1 | DeoR | WP_011002060.1 | 60 |
| OC[C@H](O)[C@@H](O)[C@H](O)C(=O)CO | L-Sorbose | WP_012490964.1 (AAF24129.1) | DeoR | AAF24132.1 | 55 |
| C=CC1=CC=C(C=C1)O | 4-vinylphenol | WP_011102052 | LysR | WP_011102053.1 | 80 |
| CCOC1=CC=CC=C1O | Guaethol | WP_027936137.1 | AraC | WP_020419855.1 | 95 |
| COc1cc(\C=C\C)ccc1O | Isoeugenol | ACP17972.1 | AraC | ACP17973.1 | 75 |
| C1=CC(=C(C=C1CCN)O)O | Dopamine | RDC20415.1 | LuxR | RDC20416.1 | 40 |
| C[N+](C)(C)CC(CC(=O)[O-])O | Carnitine | WP_003114442.1 (NP_254076.1) | AraC | NP_254073.1 | 60 |

**Supplementary Table 2:** Search terms and biosensor candidates returned by Ligify

| | CpLacI (WP_012544132.1) | PsLacI (WP_010953404.1) | TsLacI (WP_009051948.1) | LcLacI (WP_011834762.1) |
|---|---|---|---|---|
| CpLacI (WP_012544132.1) | -------------------------- | Identity: 36.13%<br>Coverage: 96% | Identity: 52.41%<br>Coverage: 98% | Identity: 29.73%<br>Coverage: 98% |
| PsLacI (WP_010953404.1) | -------------------------- | -------------------------- | Identity: 36.34%<br>Coverage: 96% | Identity: 32.90%<br>Coverage: 91% |
| TsLacI (WP_009051948.1) | -------------------------- | -------------------------- | -------------------------- | Identity: 36.23%<br>Coverage: 99% |
| LcLacI (WP_011834762.1) | -------------------------- | -------------------------- | -------------------------- | -------------------------- |

**Supplementary Table 3:** Sequence similarity of predicted D-Ribose responsive regulators.

| Alias | Metadata | | Operon and problem description |
|---|---|---|---|
| MdoR | Regulator ID: | ACS29497.1 | *No operon info* |
| | Enzyme ID: | C5MRT8 | Chemical-associate enzyme is proximal and annotated, but genome context could not be retrieved |
| | Ligand: | Methanol | |
| CamR | Regulator ID: | BAA03510.1 | *No operon info* |
| | Enzyme ID: | P00183 | Chemical-associate enzyme is proximal and annotated, but genome context could not be retrieved |
| | Ligand: | Camphor | |
| CitO | Regulator ID: | WP_002397724.1 (ENT_00500) | 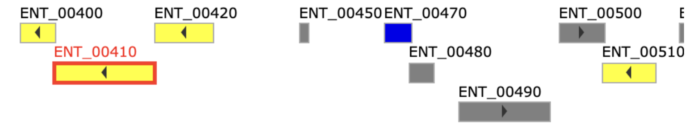  |
| | Enzyme ID: | A0A1J6XJT5 (ENT_00410) | Issue with how the operon is calculated. Doesn't include the second divergently transcribed gene |
| | Ligand: | Citrate | |
| CtcS | Regulator ID: | WP_051824537.1 (B6264_18510) | 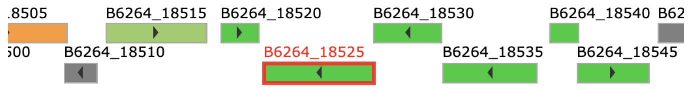  |
| | Enzyme ID: | A0A1E7MYN1 (B6264_18525) | Issue with how the operon is calculated. Doesn't include the second divergently transcribed gene. |
| | Ligand: | Chlortetracycline | |
| ClcR | Regulator ID: | AAA25771.1 (DR64_3291) | 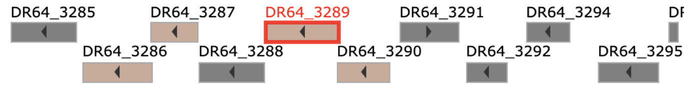  |
| | Enzyme ID: | Q13VS2 (DR64_3289) | Associated enzyme acts on 3-chloromuconate but the regulator responds to 2-chloromuconate. |
| | Ligand: | 2-chloromuconate | |

**Supplementary Figure 4**: Regulators missed due to the operon search method
